# Pangenome of white lupin provides insights into the diversity of the species

**DOI:** 10.1101/2020.06.21.163378

**Authors:** Bárbara Hufnagel, Alexandre Soriano, Jemma Taylor, Fanchon Divol, Magdalena Kroc, Heather Sanders, Likawent Yeheyis, Matthew Nelson, Benjamin Péret

## Abstract

**Background:** White lupin is an old crop with renewed interest due to its seed high protein content and high nutritional value. Despite a long domestication history in the Mediterranean basin, modern breeding efforts have been fairly scarce. Recent sequencing of its genome has provided tools for further description of genetic resources but detailed characterization is still missing.

**Results:** Here, we report the genome sequencing of several accessions that were used to establish a white lupin pangenome. We defined core genes that are present in all individuals and variable genes that are absent in some and may represent a gene pool for stress adaptation. We believe that the identification of novel genes, together with a more comprehensive reference sequence, represents a significant improvement of the white lupin genetic resources. As an example, we used this pangenome to identify selection footprints and to provide a candidate gene for one of the main QTLs associated with late flowering in Ethiopian lupin types. A 686 nucleotide deletion was identified in exon 3 of the *LaFTa1* (*Lupinus albus Flowering Time a1*) gene that suggests a molecular origin for this trait of importance, defining the need for vernalization in some lupins.

**Conclusions:** The white lupin pangenome provides a novel genetic resource to better understand how domestication has shaped the genomic variability amongst this crop. It will be of major importance for breeders to select new breeding traits and incorporate them into new, more efficient and robust cultivars in order to face a growing demand for plant protein sources, notably in Europe.

## BACKGROUND

White lupin (*Lupinus albus* L.) is a pulse whose domestication started about 3000 - 4000 years ago in the Mediterranean region [1]. It is cultivated for its seeds that contain high levels of proteins and are used both for food and feed [2]. The wild forms of the species can only be found in the Balkan region and evidence of its earliest use as a green manure and grain crop come from that same region [3]. Early Greek farmers selected larger seeds and white flowers, and presumably soft-seededness (water permeable seeds) was the earliest domestication trait. Greek and Roman literature suggests that seed indehiscence (*i.e.* resistance to pod shattering) had not yet been incorporated by the first century A.D. [4].

Wild collections and landraces of white lupin contain high levels of quinolizidine alkaloids that accumulate in the seed, resulting in a bitter taste and possible toxicity. Lysine-derived alkaloids are characteristic of the Genistoids [5–7], a monophyletic basal clade belonging to the Fabaceae family. Traditionally these bitter compounds are removed from white lupin seeds by soaking in water, a practice that is still carried out today across the Mediterranean and Nile regions [1]. However, this is uneconomic on a broad-scale, which motivated the identification of low alkaloid mutants in Germany in the 1930s, aided by advances in chemistry [4]. Modern cultivars of white lupin incorporate low alkaloid genes, hence the term ‘sweet’ lupins.

Breeding efforts have rarely been intensive or sustained over long periods. As a result, white lupin yields remain low and highly variable, in comparison to similar pulses like soybean for which important breeding efforts have been made internationally. Although white lupin cultivation represents a promising crop for Europe, in a political context aiming towards plant protein independence, the lack of well characterised genetic resources has been hampering a fast deployment of white lupin as an alternative crop to soybean imports. The recent sequencing of white lupin genome [8,9] demonstrated a resurgence of interest for this “old” crop. We believe that white lupin intragenomic diversity might reflect the early traces of its slow and sporadic domestication history.

Here we report a pangenome for white lupin that reveals important aspects of the species diversity, single nucleotide polymorphisms (SNPs) and gene presence– absence variations (PAVs). We construct a species pangenome consisting of ‘core’ genes that are present in all individuals and ‘variable’ (soft-core or shell) genes that are absent in some individuals [10,11]. Building on this comprehensive dataset, we were able to identify a deletion in the QTL region associated with late flowering in Ethiopian white lupins. The deleted gene is a homolog of the FT (Flowering Time) gene, suggesting that this deletion is at the origin of the need for vernalization in these accessions. Our analyses provide new perspectives on white lupin intra-species diversity and domestication history.

## RESULTS

### *De novo* assembly and pangenome construction

We gathered a set of 39 white lupin accessions, including 25 modern cultivars, 10 landraces and 4 wild accessions from 17 countries (Supplementary Table 1). Genome sequence of 15 out of these accessions was available from a previous report [8], whereas 24 accessions have been sequenced within this study to obtain broader species representation. Short-read sequences have been assembled *de novo* for each accession (28.5x mean depth, 150 bp pair-end, Supplementary Table 2).

The *de novo* assembly for each accession produced a total of 14.9 Gb of contigs longer than 500 base pairs (bp) with an N50 value (the minimum contig length needed to cover 50% of the assembly) of 24,475 bp. These *de novo* assemblies showed a mean complete BUSCOs score of 96.3%, a value similar to the AMIGA reference genome (97.7%). Assembly completeness assessed by BUSCO was higher than 91.7%, for all accessions and in case of three accessions (Kiev, P27174 and Magnus) the score was similar to the reference genome (Fig. 1a).

**Figure 1.**
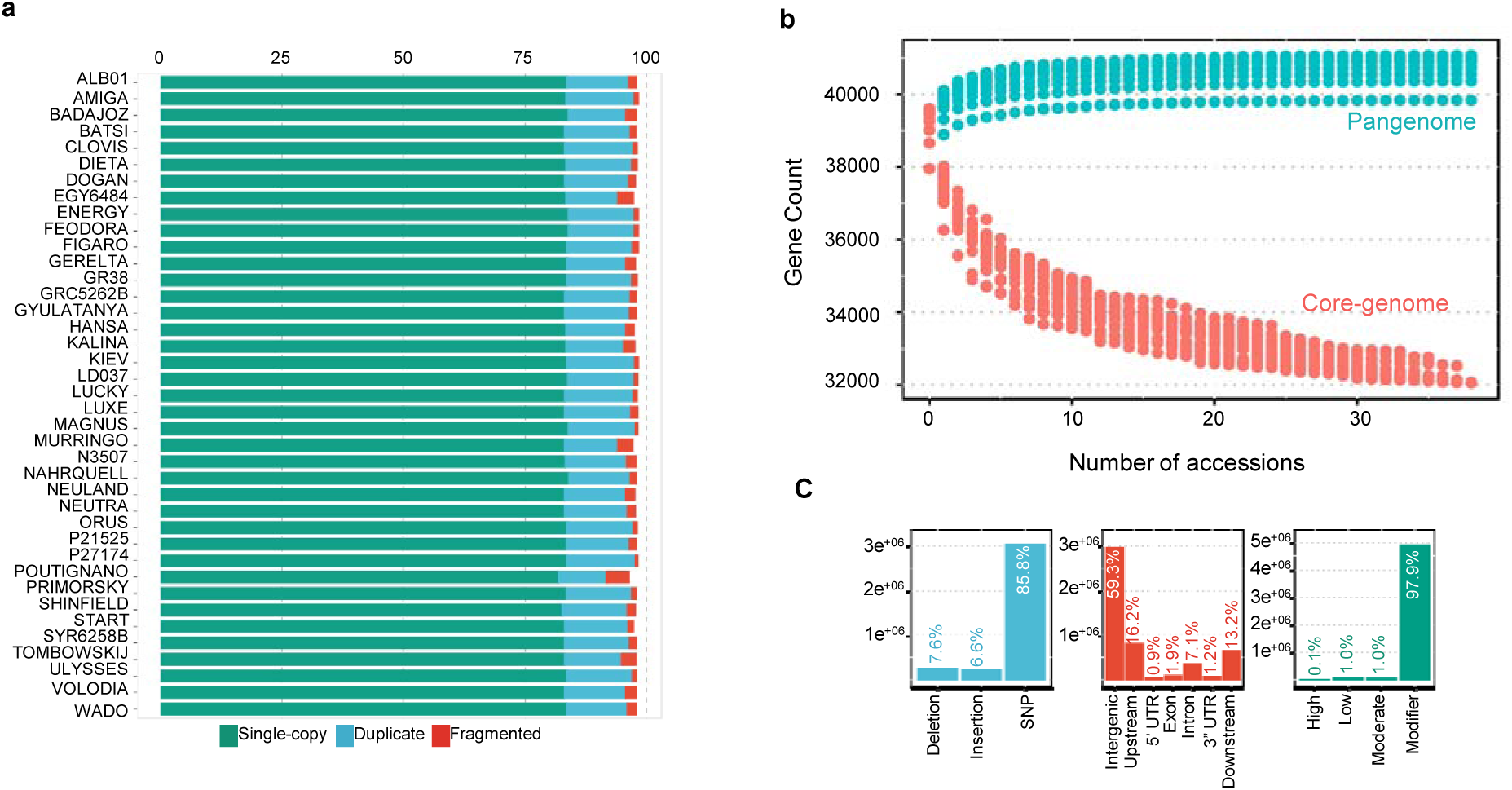
Pangenome of *L. albus*. **(a)** BUSCO percent completeness of all assemblies. All of the assemblies of this study have BUSCO completeness higher than 91.7%. **(b)** Pangenome modeling **(c)** Distribution of variants along white lupin pangenome. Types of variations identified (blue); positioning of the variants in the genome in relation to the gene structures (red); impact of the variants (green).

The pangenome was built using the iterative mapping and assembly approach, in a similar strategy used to generate the *Brassica oleracea* [10] and tomato [12] pangenomes. The assembly of *L. albus* reference genome based on AMIGA accession is 450,972,408 bp size with 38,258 predicted protein-coding genes [8]. All *de novo* assembled contigs were compared with the reference genome to identify previously unknown sequences. A total of 270 Mb of nonreference sequence with identity <90% to the reference genome was obtained. After pangenome construction and removal of contaminants and overly repetitive sequences, we assembled an additional 3,663 scaffolds, with a length greater than 2,000 bp, for a total length of 11,733,253 bp. Using a threshold of a minimum 10x coverage, we identified 178 newly predicted protein-coding genes, among which 61 could be annotated with gene ontology (GO) terms or Pfam domains (Supplementary Dataset 1). The white lupin pangenome, including reference and nonreference genome sequences, had a total size of 462,705,661 bp and contained 38,446 protein-coding genes. The total size of the constructed pangenome is compatible with nuclear DNA content estimates based on flow cytometry [13] which suggests that it represents the complete genome sequence of the species. We added to the White Lupin Genome portal (www.whitelupin.fr) dedicated user-friendly tools for the exploitation of the pangenome, such as a BLAST tool for individual accessions, download of specific regions of accessions and a genome browser mapping all the variants.

### Core and variable genes

The presence or absence of each protein-coding gene was predicted for each of the 39 accessions based on the mapping of reads from each accession to the pangenome assembly using SGSGeneLoss [14]. Likewise to other plants pangenomes [10,12,15–18], we categorized genes in the white lupin pangenome according to their presence frequencies, using Markov clustering in the GET_HOMOLOGUES-EST pipeline [19]. The majority of the genes, 32,068 (78.5%), are core genes shared by all the 39 accessions; 6,046 soft-core (14.8%), being absent in at least one accession; and 8,776 (21.4%) are shell, present in 2-38 accessions (Fig. 1b). The size of the pangenome expanded with each additional accession to 38,443 genes, and extrapolation leads to a predicted pangenome size of 40,844 +/-289 genes (Figure 1b).

### Single-nucleotide polymorphism detection and annotation

To capture and broadly characterize white lupin diversity we applied a strict SNP identification pipeline, using GATK 4.1.0.0. A total of 9,442,876 raw SNPs were identified, 806,740 of which were recognized in the newly assembled pangenome scaffolds. After filtering, 3,527,872 SNPs were retained in the 39 accessions, corresponding to a rate of 1 variant every 127 bp (Supplementary Figure S1). The majority (85.8%) of the high-quality variants are SNPs (3,027,761) and the other 501,111 variants detected are insertions and deletions (Fig. 1c – blue). Most variants (59.3%) are distributed on intergenic regions, 7.1% are within introns and only 1.9% (96,576) of the variants are located in exons (Fig. 1c – red). From the variants present in the CDS region 4,725 showed potentially large effects by causing start codon changes, premature stop codons or elongated transcripts, and 50,478 are considered to produce a moderate effect by leading to codon changes in annotated genes. The frequency of these missense SNPs in the core gene set was one each 4.26 kb, which was lower than the variable gene set, with a rate of one for 1.84 kb. The rest of the variants lead to synonymous changes in proteins (low effect variants) or modifiers, causing changes outside the coding regions (Fig. 1c – green). Collectively, this comprehensive dataset of the genome variation of white lupin provides a resource for biology and breeding of this species.

### Population structure

To establish a phylogenetic benchmark for the analysis of the pangenome, we built a consensus maximum likelihood tree (Fig. 2a) to infer the phylogenetic relationships for these *L. albus* accessions using the complete set of 3.5 M SNPs described above. This phylogenetic tree clustering supported six clades, which exhibited distinctive geographic origin and distinctive botanical features. In the Type 1 are grouped accessions with early flowering traits, including the Chilean agrogeotypes, and German and French accessions used in breeding programs. This group also included the widely used cv. Kiev Mutant, which was generated by mutagenesis techniques with the intention to induce early flowering, and the accessions that are derived forms of it (Primorsky and Dieta, [3]). Type 2 is also composed by accessions with early flowering, anumber of which have characteristics of Polish agroecotypes described by Kurlovich [3] and are adapted to grow in Eastern Europe. One of the most representative accessions of this group is the cv. Kalina [3], an old cultivar created in the Polish breeding program sharing similar genetic background with the broadly used cultivar Start. Interestingly, Start is reported to carry different early-flowering genes than Kiev Mutant [20]. Type 2 also comprises two landraces with from Syria and Israel/Palestine. Type 3 encompasses autumn-sown genotypes with strong vernalisation requirement and dwarf phenotype from the French breeding program, and the Algerian landrace ALB01. Algerian landraces are also reported to have a strong need of vernalization [3]. Type 4 comprises landraces from Iberian and Apennine Peninsula together with the described thermoneutral cultivars (*i.e.* Neutra, [2]). Type 5 is composed only by Ethiopian landraces and Wild group is composed by the four “*graecus*-type” accessions of the panel, all presenting small black-speckled seeds and non-domesticated traits (hard seeds and shattering pods).

**Figure 2.**
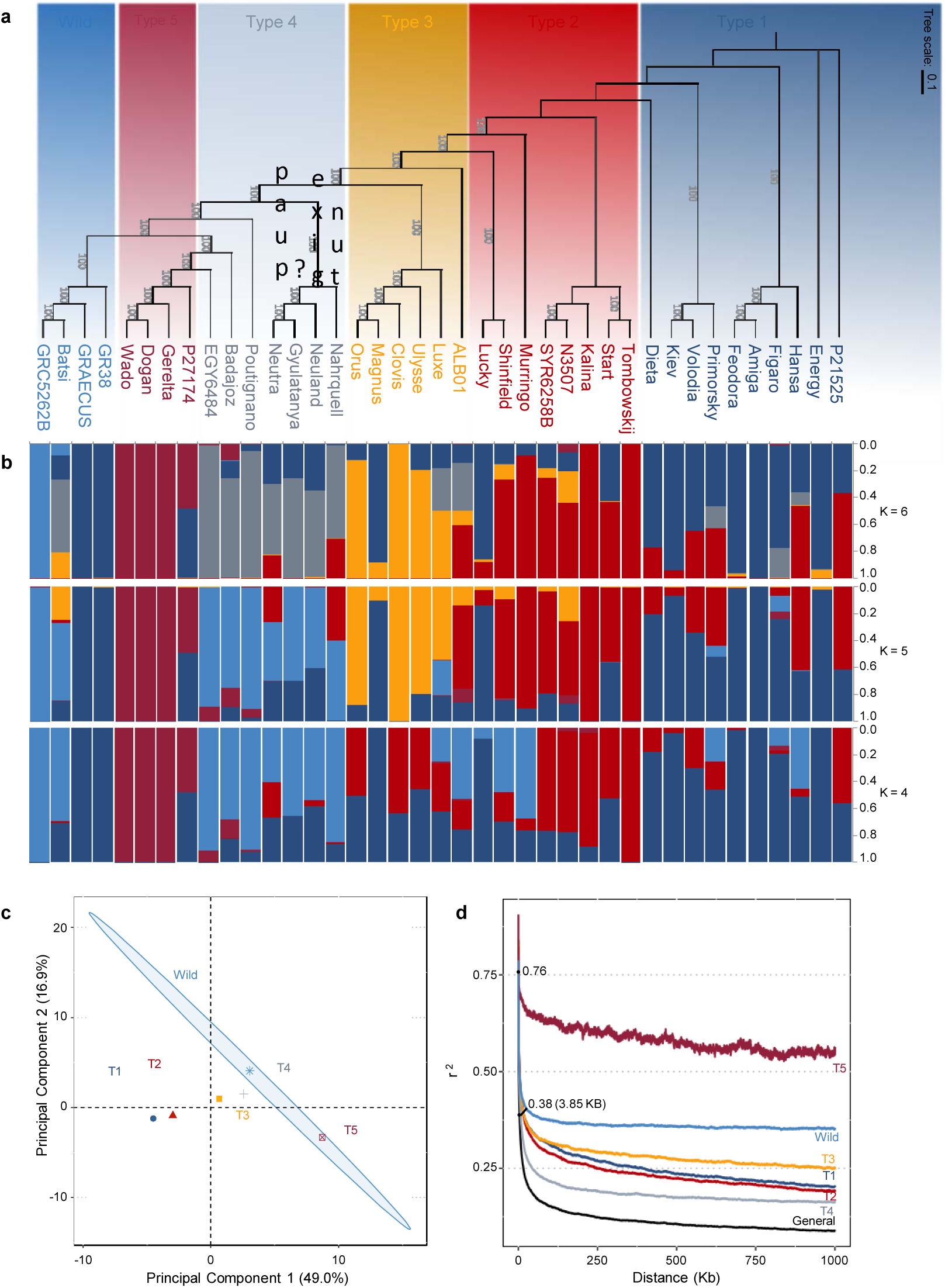
Phylogeny and population structure of 39 accessions of *L. albus*. **(a)** Maximum likelihood phylogenetic tree of white lupin constructed based on 3.5 M SNPs. The accessions are divided in 6 idiotypes. **(b)** Model-based clustering analysis with different numbers of ancestral kinships (k=4, 5 and 6). The y axis quantifies cluster membership and the x axis list the different accessions. The positions of these accessions on the x axis are consistent with those in the phylogenetic tree. **(c)** Principal component analysis based on 3.5 M SNPs. The ellipses are discriminating the accessions of each idiotype groups. **(d)** Genome-wide average LD decay estimated from different white lupin group. The decay of LD with physical distance between SNPs to half of the maximum values occurred at 3.85 kb (r^2^ =0.38) considering all accessions.

We examined genetic structure by performing a Bayesian model-based clustering analysis and found that the six population groups matched the maximum-likelihood tree (Fig. 2b). This presented evidence of significant admixture in some lines and a weak population structure, a pattern already seen in other studies of *L. albus* [21]. This weak population structure is also seen through the population-differentiation statistic (F_ST_). The F_ST_ value between all six groups were 0.27, however, F_ST_ between Type 1 and Type 2 are low as 0.086, and Type 4 and Wild have an F_ST_ of 0.092. Indeed, regarding the Bayesian model, in scenarios dividing the accessions in 4 or 5 sub-populations (Fig 2b, K=4 and K=5), accessions from Type 4 are merged with the Wild group. On the other hand, Type 5 showed a strong differentiation from the other groups, with F_ST_ values ranging from 0.34 to 0.46, with Type 4 and Type 3, respectively, which is corroborating with previous studies [22]. Principal component analysis reinforced the similarity among some groups (Fig. 2c). The two first principal components explain 65.9% of genotypic variance and it is highlighting the overlap among certain groups, in particular, Type 1 and Type 2.

Differentiation of genetic diversity between the 6 groups was investigated further through analysis of decay of linkage disequilibrium (LD, Fig. 2d). The decay of LD with physical distance between SNPs to half of the maximum values occurred at 3.85 Kb (r^2^ = 0.38), consistent with a high level of diversity and partially outcrossing mode of reproduction in this species [23]. Type 4 group also showed a fast LD decay of 5.7 Kb (r^2^ = 0.40) and Type 1-3 groups have an average LD decay of 10.5 Kb. Wild group showed a slower LD decay (38.1 Kb, r^2^ = 0.39) when comparing with the other white lupin groups, presumably an effect of the small number of wild accessions in the analysis. Nevertheless, these LD decay levels can still be considered fast compared with other plant species, for example rice (∼75–150 Kb, [24]), soybean (∼340-580 Kb, [25]) or wheat (∼7-12.4 Mb, [26]), also self-pollinated crops. The Type 5 group (Ethiopian landraces) only reached half of its LD decay after 1.5 Mb, reinforcing the high similarity of its accessions and a possible genetic isolation of this group [21]. The average nucleotide diversity π per site [27] showed that diversity was five times lower in Type 5 group (π = 0.068) compared to the general nucleotide diversity (π = 0.372). While the Wild group, although is also composed of only four accessions, showed a nucleotide diversity π = 0.402.

### Protein-coding genes presence and absence characterization

Presence and absence variants (PAVs) are an important type of structural variation and play an important role in shaping genomes, therefore contributing to phenotypic diversity [28]. The construction of a white lupin pangenome allowed identification of 1195 PAVs, representing protein-coding genes that are absent in at least one of the accessions, being 1132 genes from the reference genome and 63 from the newly identified genes (Supplementary Dataset 2-3). We further examined if the phylogenetic groups have an influence in the number of PAVs and if the PAVs are homogeneous within the groups (Fig. 3a-c). The wild accessions have a significatively higher number of newly identified genes, with the accessions GRAECUS and GR38 only missing 4 of them. The four wild accessions share 157 out of the 178 new-identified genes in the pangenome (Fig. 3a).

**Figure 3.**
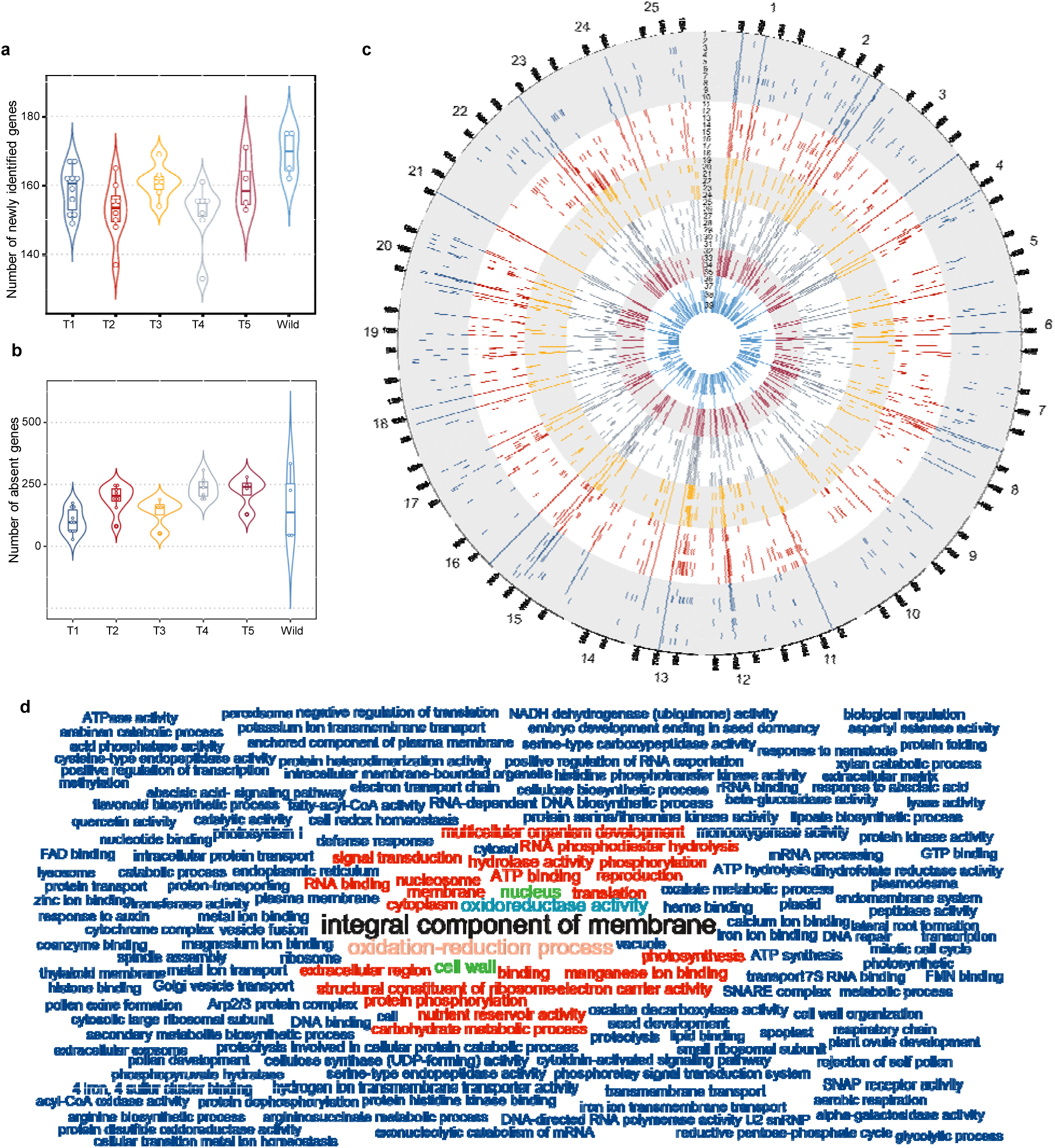
PAV of coding gene in *L. albus*. **(a)** Number of newly identified genes by phylogenetic group. **(b)** Number of absent genes by phylogenetic groups. **(c)** Positioning of absent genes in the 25 white lupin chromosomes in each one of the 39 accessions. Order of accessions from outer to inner track: 1-AMIGA, 2-FEODORA, 3-FIGARO, 4-ENERGY, 5-KIEV MUTANT, 6-HANSA, 7-P21525, 8-PRIMORSKY, 9-DIETA, 10-VOLODIA, 11-START, 12-N3507, 13-TOMBOWSKIJ, 14-KALINA, 15-SYR6258B, 16-LUCKY, 17-MURRINGO, 18-SHINFIELD, 19-ALB01, 20-LUXE, 21-ULYSSE, 22-MAGNUS, 23-CLOVIS, 24-ORUS, 25-NAHRQUELL, 26-GYUNLATANYA, 27-NEULAND, 28-NEUTRA, 29-BADAJOZ, 30-EGY6484B, 31-POUTIGANO, 32-P27174, 33-GERELTA, 34-DOGAN, 35-WADO, 36-GR38, 37-GRAECUS, 38-BATSI, 39-GRC5262B. The accessions’ colors reflect the 6 idiotypes. **(d)** Functional enrichment analysis of the variable genome. Graphical representation of enriched biological process (GOs). Size of the words and colors are proportional to their representativeness in the gene pool.

The number of missing genes within individual genomes ranges from 45 (AMIGA – Type 1) to 348 genes (GRC5262B – Wild). Each group shares a median of 31 common lost genes amongst all its accessions and a total of 103 genes are absent in at least one accession of each group (Fig. 3b). There are 137 genes that have been exclusively lost within accessions of the Wild group, however only 30 genes are shared among all the *graecus* accessions. On the other hand, genomes of Ethiopian landraces (Type 5) share a total of 118 common missing genes, amongst 39 are unique for this group. Remarkably, for this group there is a concentration of lost genes on Chr17. This includes a set of 9 tandem duplicated genes covering a region of 120 Kb (Supplementary Fig. 2). They are annotated as “Putative ferric-chelate reductase (NADH)” homologs of *Arabidopsis* gene *FRO2*, known for its role of iron uptake by the roots under stress condition [29].

Checking the position of the PAVs on the chromosomes we could identify some peculiarity regarding the PAVs within the groups. For example, on Chr13 there is a concentration of PAVs in the region of 5-10 Mb that are missing from most accessions of Type 2-5 and Wild, but are present in the genomes of most Type 1 members. Similar pattern happens in the 3.6-6.4 Mb region of Chr04. Chr23 has the highest number of PAVs (78), a common feature of all the groups.

Functional analysis of PAVs suggests enrichment of GO terms as “integral component of membrane” (GO:0016021) and “oxidation-reduction process” (GO:0055114) (Fig. 3d, supplementary Fig. 3 and Supplementary Data 2). These suggest an enrichment of genes and gene families coding for membrane receptors proteins or membrane transporters. Other GO terms suggest that some of the genes may be involved in cell wall remodeling (“cell wall” - GO:0005618; “cell wall organization” - GO:0071555). Genes with these functions are frequently linked to biotic and abiotic stress responses [30,31]. PAV genes related to abiotic and biotic stress responses have been observed in several plant species [15,17,18,32–35] and these may reflect the evolution for adaptive traits related for each agroecotype. Moreover, the presence/absence of these stress-response related genes may also be partially due to whole-genome triplication event on white lupin genome [8], which caused an overlapping roles in various loci.

### Footprints of selection and alleles identification in candidate genes

To demonstrate the power of white lupin pangenome to address basic research questions, we used it to detect possible footprints of selection and to identify alleles in candidate genes underlying major QTLs. Firstly, to examine potential selective signals during white lupin domestication and breeding, we scanned white lupin genome searching for regions with marked reductions in nucleotide diversity (Fig. 4).

**Figure 4.**
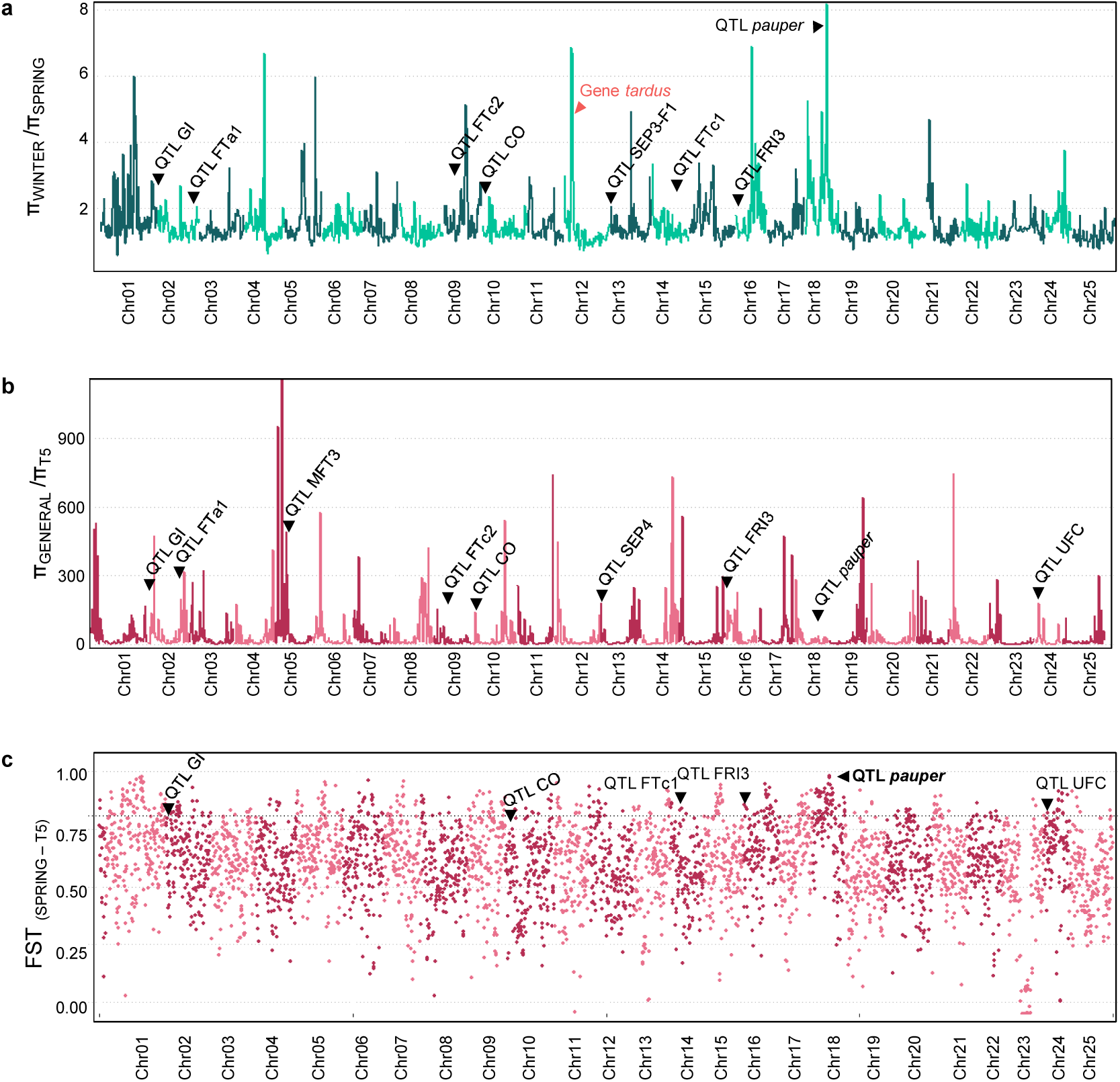
Footprints of selection in the white lupin genome. Nucleotide diversity (π) comparison between **(a)** Winter (T3 and T4) and Spring accessions (T1 and T2) and **(b)** between all accessions (General) and Ethiopian accessions (T5). QTLs previously reported and a *L. angustifolius* domestication gene (red) that overlapped with selective sweeps are marked. **(c)** Fst-based genome-wide analysis of population differentiation estimated between Spring (T1 and T2) and Ethiopian (T5) accessions. Black horizontal dashed line marks the .90 percentile of distribution of Fst estimated (Fst = 0.81).

The domestication and breeding efforts in white lupin have focused in searching for accessions with reduced seed alkaloid content, reduced time to flower as well as excessive indeterminate branching. Therefore, we combined Type 1 and Type 2 accessions, that are spring types and went to a more intense breeding process and compared them with Type 3 and Type 4 accessions, that are winter types (Fig. 4a). A selective sweep affecting only the spring white lupin accessions would be expected to leave a typical low-polymorphism and high-divergence signal around the region of the selected genes. We measured the sweep on the nucleotide diversity (π value [27]), by comparing the two groups (πWinter/πSpring) over 250-kb windows. We identified 167 putative selection sweeps associated to the breeding of the spring accessions (πWinter/πSpring > 2.101). We observed that some of the peaks co-localized with previously reported white lupin QTLs for flowering time and alkaloid content [36,37]. The same pattern was observed when checking for the divergence of the gene pool between these two groups along the chromosomes (Fst, Supplementary Fig. 4a).

Interestingly, other peaks with higher sweeps of diversity are present, indicating that other genomic regions may be implicated with these traits and may carry other important genes of these pathways. Furthermore, they highlight specific genomic regions of spring accessions that have been selected during domestication and breeding. For instance, we checked for orthologs of domestication genes from the close relative narrow-leafed lupin and found that the gene Lalb_Chr12g0203121, a homolog of a candidate gene for the reduced pod shattering locus *tardus* (Lup002448, [38]), is co-localized with a sweep peak on Chr12.

The reported white lupin QTLs were identified in a recombinant inbred line (RIL) mapping population derived from the cross between Kiev Mutant (Type 1) and the Ethiopian landrace P27174 (Type 5). Thereupon, we checked the sweep of diversity between all accessions compared to Ethiopian accessions, T5 (πGeneral/πT5, Fig. 4b) and identified 84 sweep peaks (πGeneral/πT5 > 83.97). A similar trend of co-localization of the QTL peaks were observed, with steep peaks around the QTL regions. Interestingly, the region corresponding the QTL *pauper* did not show a peak, being far below the statistical significance threshold. This indicates that the two groups have similar level of nucleotide diversity in this region (Supplementary Fig. 4b). It can be explained by the above-mentioned similarities among accessions of group T5 and that many of the modern accessions carry the low alkaloid alleles for the pauper region. However, although this region showed a similar nucleotide diversity between the two groups, it presented a high genetic variance, with a median FST of 0.94 for the region (Fig. 4c).

In another approach to demonstrate the power of white lupin pangenome, we used its assembly to identify a candidate gene underlying a major QTL and describe the associated allelic diversity. Chromosome 2 is the location of an important QTL associated with early flowering white lupins. We used the protein sequences of *Lupinus angustifolius* that have been previously mapped in syntenic regions of the these QTLs [39] to perform an homology search against the pangenome. we identified the gene *LaFTa1* (Lalb_Chr02g0156991), a homolog of the gene *LanFTa1* (Lup21189) mapped on this QTL region. The white lupin *LaFTa1* (Lalb_Chr02g0156991) was annotated as “Putative phosphatidylethanolamine-binding protein” (PEBP) in the reference genome. The FT proteins belonging to the PEBP family are the key control points of the flowering time in plants. The *LaFTa1* gene presented a deletion of 686 on the third intron that is present only on Type 5 accessions, that have late flowering phenotypes (Fig. 5a-b and Supplementary Fig. 5). Indeed, one of the parents of this QTL mapping population belongs to this group (P27174). It is reported that changes in FT promoter and introns can alter FT expression in response to photoperiod and vernalization, and consequently, induce flowering [40]. This suggests that the identified *LaFTa1* is the gene underlying this QTL and that this deletion on the intron of Type 5 accessions may be contributing for the late flowering pattern of this group.

**Figure 5.**
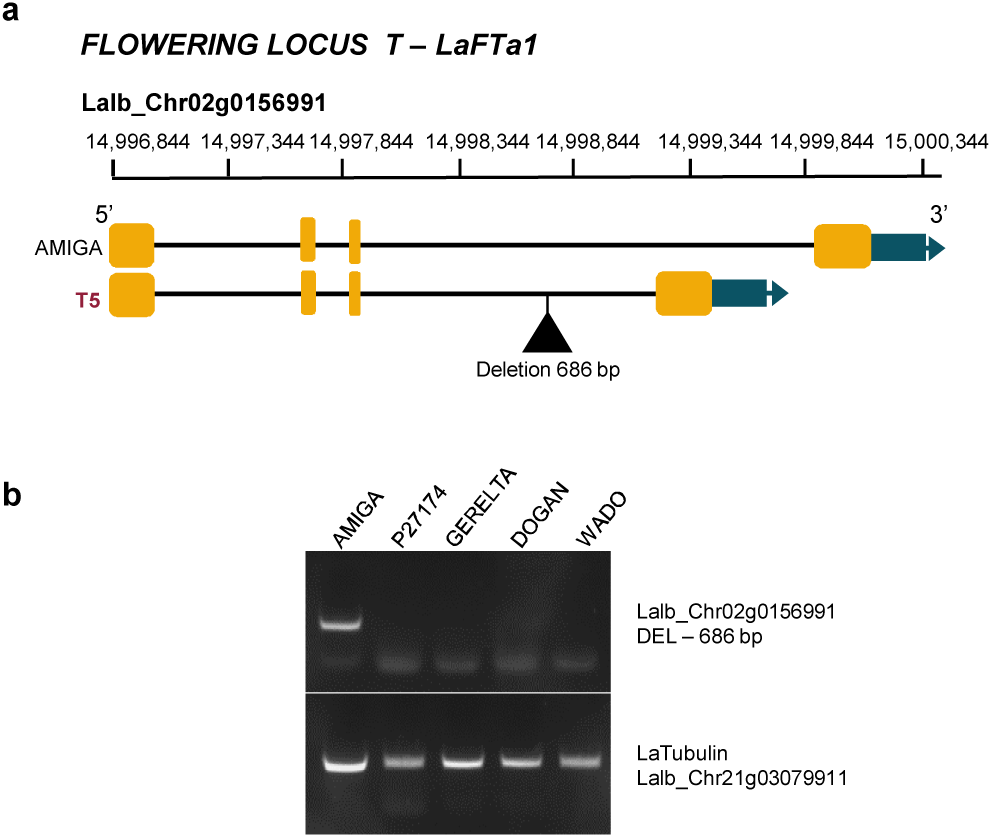
Identification and allele variation of candidate gene *FLOWERING LOCUS T.* **(a)** Candidate gene located on chromosome 2. Type 5 accessions, originated from Ethiopia, have a deletion of 686 bp in the third intron. **(b)** Confirmation of the deletion in the third intron of Type 5 accession by PCR. Gene *LaTubulin* was used as positive control.

## DISCUSSION

A pangenome is a complete set of genes for a species, including core genes which are present in all individuals, and variable genes which are absent in one or more individuals [41]. We generated a *de novo* assembly for 38 white lupin accessions and, taking advantage of a good reference assembly for the species [8], we constructed a *L. albus* pangenome by iteratively and randomly sampling these sequenced accessions. This dataset is representative of the diversity of the species, containing wild accessions, landraces and cultivars of white lupin from across their respective distributions. As a result, we estimate that this white lupin pangenome assembly effectively encompasses the complete sequence for the genome of the species, with 462,7Mb sequence and containing 38,446 protein-coding genes. The finding that 21.5% of genes in the pangenome exhibit varying degrees of genic presence/absence variants (PAVs) highlights the diverse genetic feature of white lupin and the significant improvement of the reference genome, by including genomic information of other accessions and discovery of new genes. Remarkably, the white lupin pangenome showed a high content of core genes (78.5%), as compared with other plant species as tomato (74.2%, [34]), *Arabidopsis thaliana* (70%,[19]), bread wheat (64%, [42]), sesame (58%, [16]) and wild soybean (49%, [43]), which might be a reflection of its domestication history and modest breeding efforts to date.

The domestication of white lupin started during Bronze Age [4], and the ancestral history of this species is different than other major crops such as rice, maize, sorghum, tomato, and soybeans, which are more ancient [44]. The early cultivated forms have the same Mediterranean distribution that its wild ancestor types (*graecus*), which led to small adaptation or selection differences. *L. albus* domestication was slow with potentially centuries between acquisition of each domestication trait, which may explain why there is not a more pronounced genetic differentiation between wild, landrace and cultivated types [45]. This is echoed in the lack of population structure presented within these accessions and in the low LD extent, which generally reduce the diversity and change allele frequencies either to fixation or intermediate frequencies [46]. Despite being a largely self-pollinating crop (with an out-crossing rate reported as 8–10 %, [23]), white lupin showed a remarkably low LD extent (< 4kb), even lower than the wild population of its relative, narrow-leafed lupin, that showed a decay of LD after 19.01 Kb [47]. One distinction between these closely related species is that narrow-leafed lupin is almost exclusively self-pollinating and so the modest levels of outcrossing in white lupin may be a key factor governing the differences in LD between these two species. Having a low LD and weak population structure together mean that association mapping is likely to be particularly powerful in white lupin, in contrast to the more highly structured and high LD species narrow-leafed lupin, where association studies have so far proved rather weak [47,48].

Type 5 accessions, from Ethiopia, are the only group which shown a strong genetic differentiation from the others, with F_ST_ values higher than 0.3. Such a distinct separation is an evidence that the Ethiopian accessions have evolved in isolation and the genetic differences are probably due to ancient founder effects. The differences of Type 5 group are also highlighted by the PAVs. Together with the Wild group, Ethiopian landraces carry most of the new identified genes and also miss a large number of genes of the reference genome (Fig. 3). Moreover, it is a highly homogeneous group, with all accessions sharing a large number of these lost genes. The loss of these genes is probably an adaptive response for the local environment. For instance, the loss of the nine tandem duplicated homologue *AtFRO2* on Chr17 might be an adaptive response to highland Ethiopian soils that are iron-rich [49]. A more detailed look into the PAVs among the different groups may be useful to better understand their specificities.

Our analysis brings a high resolution to the within-species diversity. Using the pangenome dataset, we performed genome-wide comparisons of the assemblies, enabling the characterization of more than 3 million complex variants, including many large-effect coding variants which should be helpful in pinpointing causal variations in QTLs for important traits and in future genome-wide association studies. In particular, our study demonstrated that 4,725 genes were found to contain important coding variation in at least one accession and might have important biological functions underlying the variation of complex traits.

We wanted to demonstrate how a pangenome can be a useful tool to identify allelic differences that are responsible for phenotypic variation. By performing a genome wide analysis, we detected that nucleotide diversity were quite variable across the genome. The efforts of breeding in white lupin have been focused in combining of domestication traits such as soft and white seeds and reduced pod-shattering, which were already available from ancient times, with that of reduced alkaloids, increased yield and the reduction of flowering time and excessive branching [45]. Looking for differences in nucleotide diversity across the genome amongst breeding accessions and comparing with landraces/wild accessions, we could detect some peaks of sweep of diversity. In these peaks there is an important decrease of nucleotide diversity within the breeding lines and they represent marks of selection (Fig. 4). In these regions were also detect a high divergence between the two gene pools (Fst). However, although the sweeps of diversity co-localize with some identified QTLs for flowering time and low alkaloid content, there are other higher peaks along the chromosomes. These regions should be explored in order to find genes underlying phenotypic traits that have been selected directly or indirectly during domestication and breeding of white lupin. For instance, white lupin is known for thriving in soils with low nutrient availability by producing specialized root structures called cluster roots [50]. In a previous work, we demonstrated that the breeding accessions have an earlier establishment of the root system through lateral and cluster root formation that was indirectly selected [8]. By looking closer in these chromosome regions with low nucleotide diversity and high genetic differentiation we might be able to find genes with important roles in the root architecture of white lupin. Hence, integration of the information from studies of gene function and the high density of variants described in this pangenome can provide a complementary approach to forward genetic studies and can contribute to develop the research and breeding of white lupin.

## CONCLUSION

In summary, the white lupin pangenome comprises a wealth of information on genetic variation that has yet to be fully exploited by researchers and breeders. Although a there is large collection of white lupin accessions available in genebanks worldwide, they barely have been explored and genetically characterized. This pangenome represents a comprehensive and important resource to facilitate the exploration of white lupin as a legume model for future functional studies and molecular breeding.

## METHODS

### Genome sequences of white lupin accessions

We retrieved the genome sequencing data of 15 white lupin accessions that were published previously [8], including 11 modern cultivars, 1 landrace and 2 wild relatives. They were sequenced using Illumina technology using paired-end 2 × 150 bp short-reads with average sequencing depth of 45.99×. It included Illumina genome data of 64.47x depth for the reference cultivar “AMIGA”. Genome sequences of additional 24 accessions were generated here, including 12 modern cultivars, 9 landraces and 2 wild relatives. Young leaves of individual plants were used to extract genomic DNA of each accession using the QIAGEN DNeasy Plant Mini kit following the supplier’s recommendations. The accessions were sequenced using Illumina technology using paired-end 2 × 150 bp short-reads (Macrogen, South Korea). It was generated a total of 196.85 Gb of data with average sequencing depth of 19.1x. (Supplementary Table 2).

### De novo genome assembly and pangenome construction

Reads were processed to trim adapters and low-quality sequences using Cutadapt 1.15 [51] with parameters ‘--pair-filter=any -q20,20 -m 35’ and the forward and reverse Illumina TruSeq Adapters. The final high-quality cleaned Illumina reads from each sample were *de novo* assembled using Spades 3.13.0 [52] with k-mer size of 21,33,55,77,99,121. The assembled contigs were then aligned to the white lupin reference genome [8] (GenBank accession no.: WOCE00000000, http://www.whitelupin.fr.), using the steps 7 and 8 of the EUPAN Pipeline [53], in order to extract contigs that were not aligning to the reference. Then, redundancy in the extracted contigs has been reduced using CD-hit 4.8.1 with default parameters. The resulting contigs were then search against the NCBI nt nucleotide database using blastn 2.10 [54]. Sequences with best hits from outside the Eudicots, or covered by known plant mitochondrial or chloroplast genomes, were possible contaminations and were therefore removed.

### Annotation of the white lupin pangenome

A custom repeat library was constructed by screening the pangenome and the white lupin reference genome using RepeatModeler (http://www.repeatmasker.org/RepeatModeler/), and used to screen the nonreference genome to identify repeat sequences using RepeatMasker (http://www.repeatmasker.org/). Contigs with more than 98% of repetitive sequences were removed from the annotation pipeline. Protein-coding genes were predicted from nonreference genome using MAKER2 [55]. *Ab initio* gene prediction was performed using Augustus [56] and SNAP [57]. Augustus [58] has been previously trained for white lupin as described in the documentation, and SNAP was trained for two rounds based on already assembled transcriptome of white lupin, as described in maker2 documentation. In addition, protein sequences of white lupin, *Medicago truncatula* and the Viridiplantae subset of Swissprot were used as evidence. Finally, gene predictions based on *ab initio* approaches, and transcript and protein evidence were integrated using the MAKER2 pipeline. A set of high-confidence gene models supported by transcript and/or protein evidence were generated by MAKER2. In order to remove possible remaining contamination, all high confidence maker generated protein sequences were aligned against the nr databses, and sequences with best hits from outside Eudicots or with best hit inside chloroplastic and mitochondrial sequences were removed. Genes that matched white lupin reference sequences were also removed the same way.

In parallel, contigs with a length superior to 2Kb from the whole assembly of the 39 lupin accession were annotated using the Egnep 1.5.1 pipeline [59]. RepeatMasker was used to detect and remove contigs constitute by more than 98% of known repeat sequences based on the previously built white lupin repetitive element sequences database. The white lupin transcriptome [8]was used as ESTs evidence, using a minimum identity percentage of 95%, along with the proteome of white lupin, *Medicago truncatula*, and the Viridiplantae subset of the swissprot database, with weight of 0.4, 0.3 and 0.3 respectively. Resulting predicted proteins were search against REXdb and repbase in order to remove possible transposable elements. The resulting genes prediction were again scan with repeat-masker, and genes composed of more than 90% of detected repetitive sequences were removed from further analyses in order to control false positive.

### Gene presence/absence variation and pangenome modeling

Reads were processed to trim adapters and low-quality sequences using Cutadapt 1.15 [51] with parameters ‘--pair-filter=any -q20,20 -m 35’ and the forward and reverse Illumina TruSeq adapters. Resulting high quality reads were then aligned to the pangenome using BWA-MEM [60] with default parameters. Picard tools was used to remove possible PCR and optical duplicates, and reads considered as not properly paired were removed using samtools view. The presence or absence of each gene in each accession was determined using SGSGeneLoss [14]. In brief, for a given gene in a given accession, if less than 10% of its exon regions were covered by at least five reads (minCov = 5, lostCutoff = 0.1), this gene was treated as absent in that accession, otherwise it was considered present. The parameters used for the new set of gene discovered in the pangenome were different: minCov = 10 and lostCutoff = 0.8. For more precise pangenome studies taking into account all the genes discovered in all the different varieties, GET_HOMOLOGUES_EST was used on the whole CDS and proteome of the whole 39 varieties with parameters “-R 123545 -P -M -c -z -A -t 2” to detect clusters of genes shared by at least two varieties.

### SNP discovery and annotation

Cutadapt [61] was used to remove Illumina Truseq adapters from the sequencing data and to remove bases with a quality score lower than 30, in both 5’ and 3’ end of the reads. Reads with a length lower than 35 were discarded. We then used BWA-MEM version 0.7.17 [60] to map the resequencing reads from all 39 genotypes to the white lupin reference genome. PCR and Optical duplicates were detected and removed using Picard Tools. After that, GATK 4 HaplotypeCaller tool was used in emit-ref-confidence GVCF mode to produce one gvcf file per sample. These files were merged using GATK Combine GVCFs. Finally, GATK GenotypeGVCFs was used to produce a vcf file containing variants from all the 39 samples. This identified a total of 9,442,876 SNPs/indel. After filtering for minimum allele frequency of 0.15 and heterozygosity frequency of 0–0.2, 3,527,872 SNPs were retained for further analysis.

### Evolutionary analysis

A maximum-likelihood phylogenetic tree was constructed based on 3,121,673 parsimony-informative SNPs with 1,000 bootstraps using IQ-TREE [62] using ModelFinder [63] option. Then, a phylogenetic tree was prepared using the iTOL v 4.3 [64].

Population structure based on the same set of SNPs was investigated using STRUCTURE [65]. Thirty independent runs for each K from 1 to 15 were performed with an admixture model at 50,000 Markov chain Monte Carlo (MCMC) iterations and a 10,000 burn-in period. Principal component analysis using this SNP dataset was performed using the function “princomp” in R (http://www.R-project.org/). The linkage disequilibrium (LD) pattern was computed using PopLDdecay v3.40 [66]. LD decay was measured on the basis of the r^2^ value and the corresponding distance between two given SNPs.

### Selective sweep analyses

To detect genomic regions affected by domestication we used the same set of 3,121,673 SNPs using Tassel [67]. The level of genetic diversity (π) was measured with a window size of 2000 SNPs and a step size of the same length, generating windows of approximately 250-kb. Genome regions affected by selection or domestication should have substantially lower diversity in spring white lupin (Types 1 and 2, πSpring) than the diversity in winter accession (Type 3 and 4, πWinter) and Ethiopian accessions (πT5). Windows with the top 10% highest ratios of πWinter/πSpring (≥2.101) or πGeneral/πCA (≥83.969) were selected as candidate selection and domestication sweeps. The PopGenome package [68] in R with its sliding window method was used to calculate the interpopulation differentiation, FST. Using a set of 40 k random good-quality SNPs evenly distributed along the 25 chromosomes, we calculated nonoverlapping sliding-windows of 10 SNPs each.

## DECLARATIONS

### Ethics approval and consent to participate

Not applicable

### Consent for publication

Not applicable

### Availability of data and materials

The detailed methods and datasets supporting the conclusions of this report are included within the article and its additional files. All deep sequencing data reported in this paper have been submitted to the NCBI. The datasets generated and analyzed during the current study are available from the corresponding author upon request. Full genomic and raw sequence data are publicly available for download on the White Lupin genome portal [www.whitelupin.fr/pangenome] that contains a Genome Browser, Expression tools and a Sequence retriever dedicated to the pangenome. The pangenome project and raw data has been deposited at DDBJ/ENA/GenBank under the accession PRJNA608889.

### Authors’ contributions

A.S. developed bioinformatic resources and performed pangenome assembly. J.T. and F.D. performed DNA extraction and experiments. M.N., H.S., L.Y. and M.K. provided genetic material. B.H. performed data analysis. B.H., M.K., M.N. and B.P. designed experiments and wrote the article.

