## Supplementary Figures for "Pangenome of white lupin provides insights into the diversity of the species"

### Slide 1
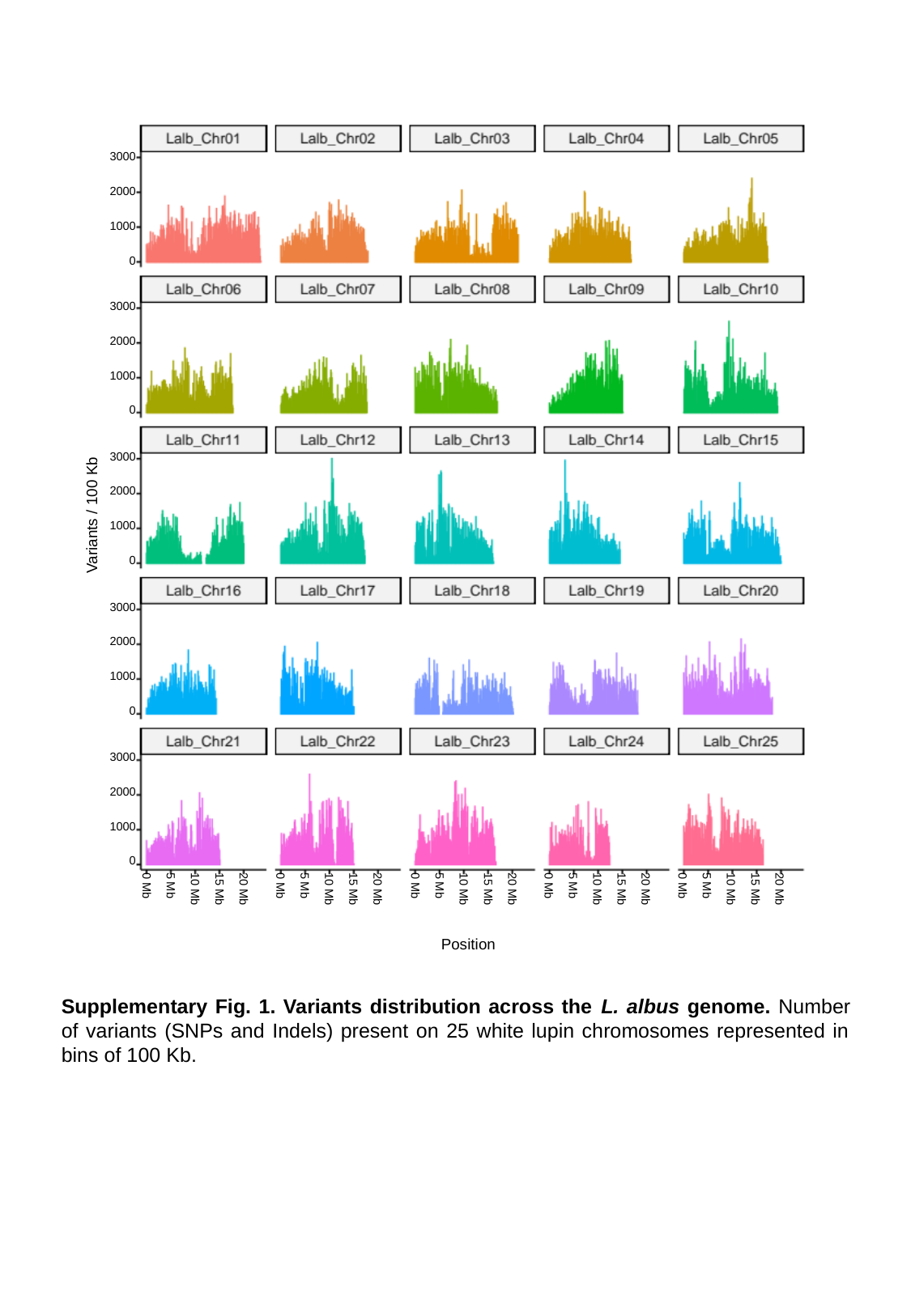

3000
2000
1000
0
3000
2000
1000
0
3000
2000
1000
0
3000
2000
1000
0
3000
2000
1000
0
0 Mb
0 Mb
0 Mb
0 Mb
0 Mb
5 Mb
5 Mb
5 Mb
5 Mb
5 Mb
10 Mb
15 Mb
20 Mb
10 Mb
15 Mb
20 Mb
10 Mb
15 Mb
20 Mb
10 Mb
15 Mb
20 Mb
10 Mb
15 Mb
20 Mb
Variants / 100 Kb
Position
Supplementary Fig. 1. Variants distribution across the L. albus genome. Number of variants (SNPs and Indels) present on 25 white lupin chromosomes represented in bins of 100 Kb.

### Slide 2
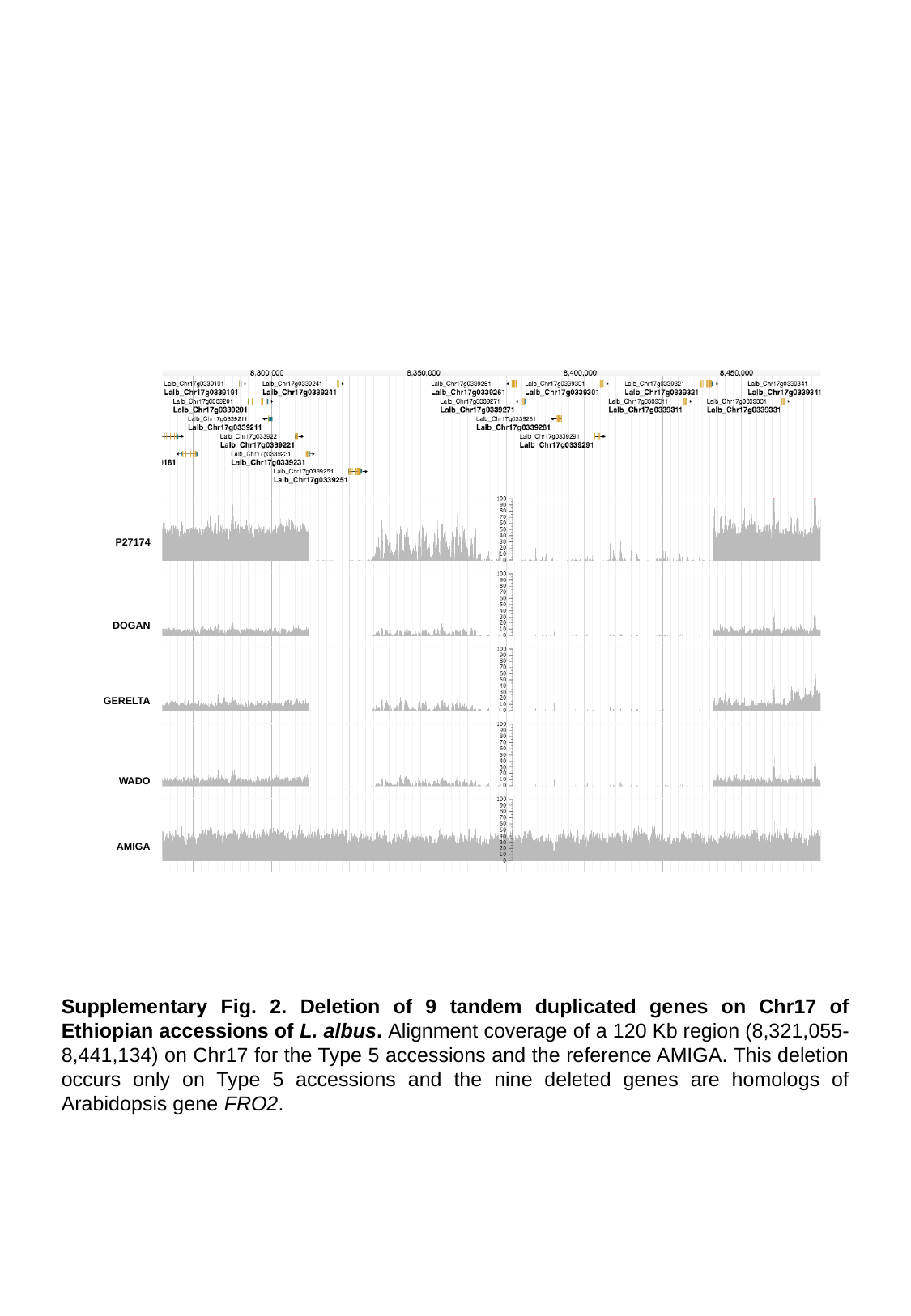

P27174
DOGAN
GERELTA
WADO
AMIGA
Supplementary Fig. 2. Deletion of 9 tandem duplicated genes on Chr17 of Ethiopian accessions of L. albus. Alignment coverage of a 120 Kb region (8,321,055-8,441,134) on Chr17 for the Type 5 accessions and the reference AMIGA. This deletion occurs only on Type 5 accessions and the nine deleted genes are homologs of Arabidopsis gene FRO2.

### Slide 3
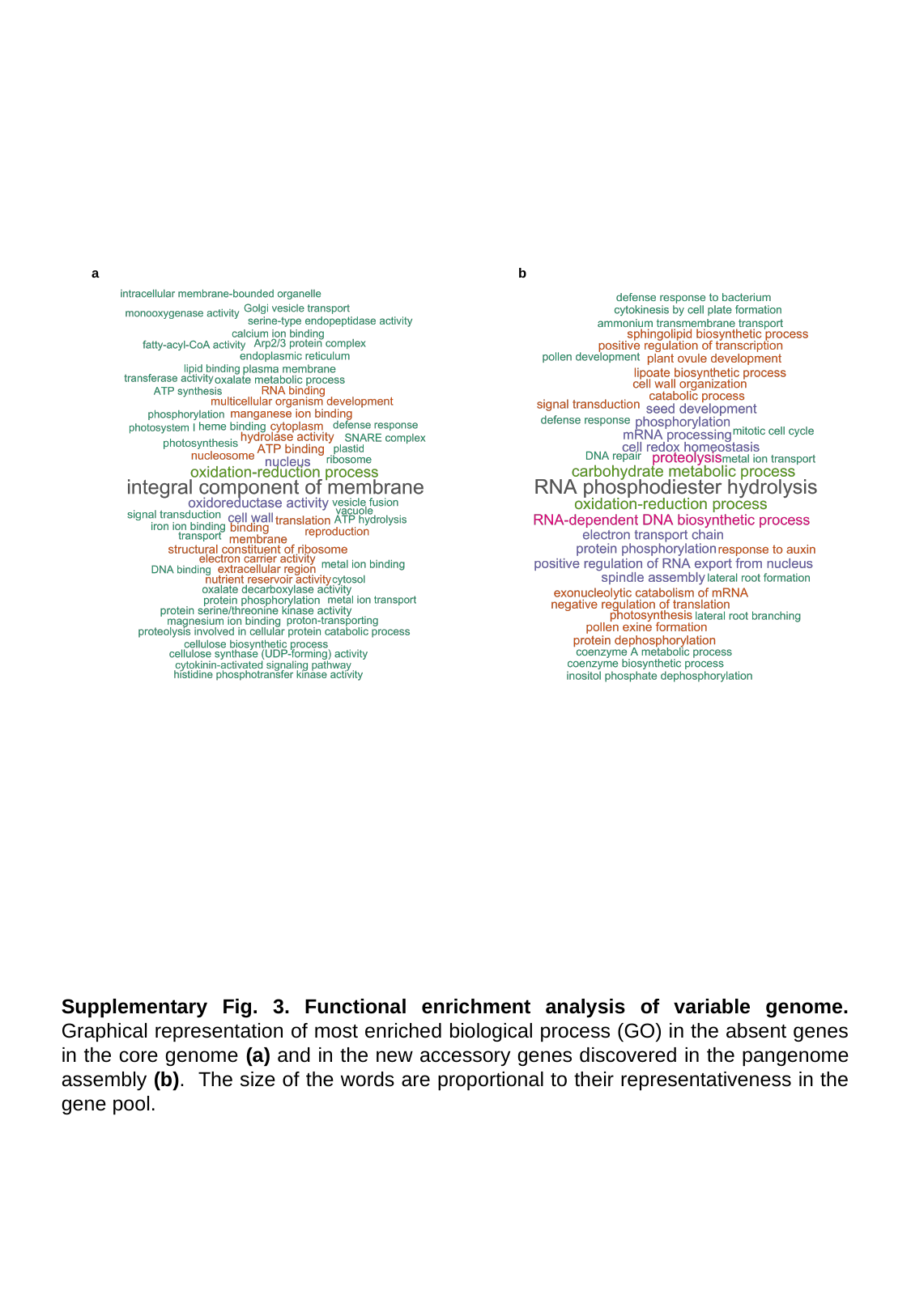

a
b
Supplementary Fig. 3. Functional enrichment analysis of variable genome. Graphical representation of most enriched biological process (GO) in the absent genes in the core genome (a) and in the new accessory genes discovered in the pangenome assembly (b). The size of the words are proportional to their representativeness in the gene pool.

### Slide 4
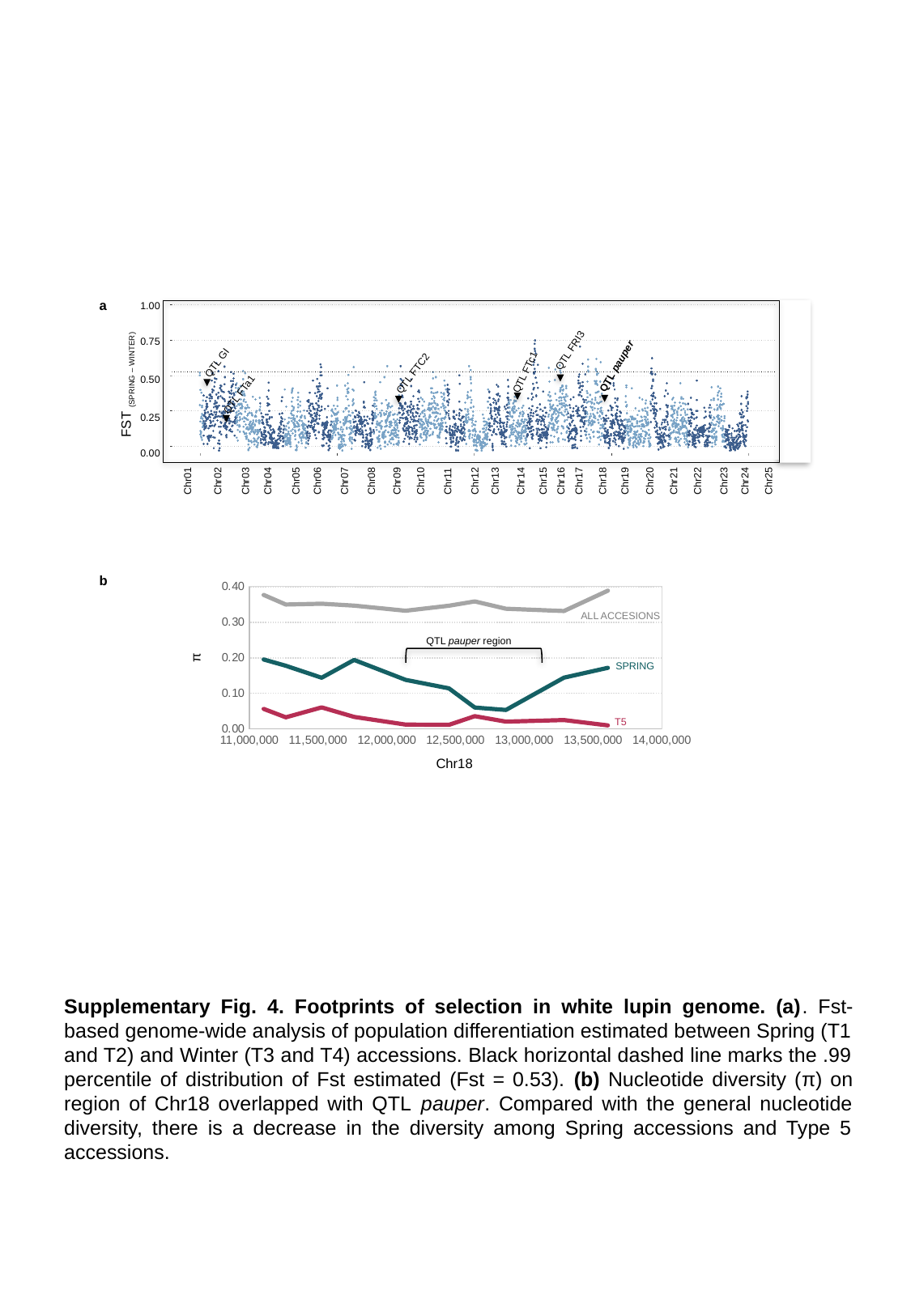

a
1.00
0.75
FST _(spring – winter)_
0.50
0.25
0.00
Chr01
Chr02
Chr03
Chr04
Chr05
Chr06
Chr07
Chr08
Chr09
Chr10
Chr11
Chr12
Chr13
Chr14
Chr15
Chr16
Chr17
Chr18
Chr19
Chr20
Chr21
Chr22
Chr23
Chr24
Chr25
QTL FRI3
QTL FTc1
QTL GI
QTL pauper
QTL FTC2
QTL FTa1
b
#### Chart
| Category | SPRING | T5 | ALL |
|---|---|---|---|
ALL ACCESIONS
QTL pauper region
π
SPRING
T5
Chr18
Supplementary Fig. 4. Footprints of selection in white lupin genome. (a). Fst-based genome-wide analysis of population differentiation estimated between Spring (T1 and T2) and Winter (T3 and T4) accessions. Black horizontal dashed line marks the .99 percentile of distribution of Fst estimated (Fst = 0.53). (b) Nucleotide diversity (π) on region of Chr18 overlapped with QTL pauper. Compared with the general nucleotide diversity, there is a decrease in the diversity among Spring accessions and Type 5 accessions.

### Slide 5
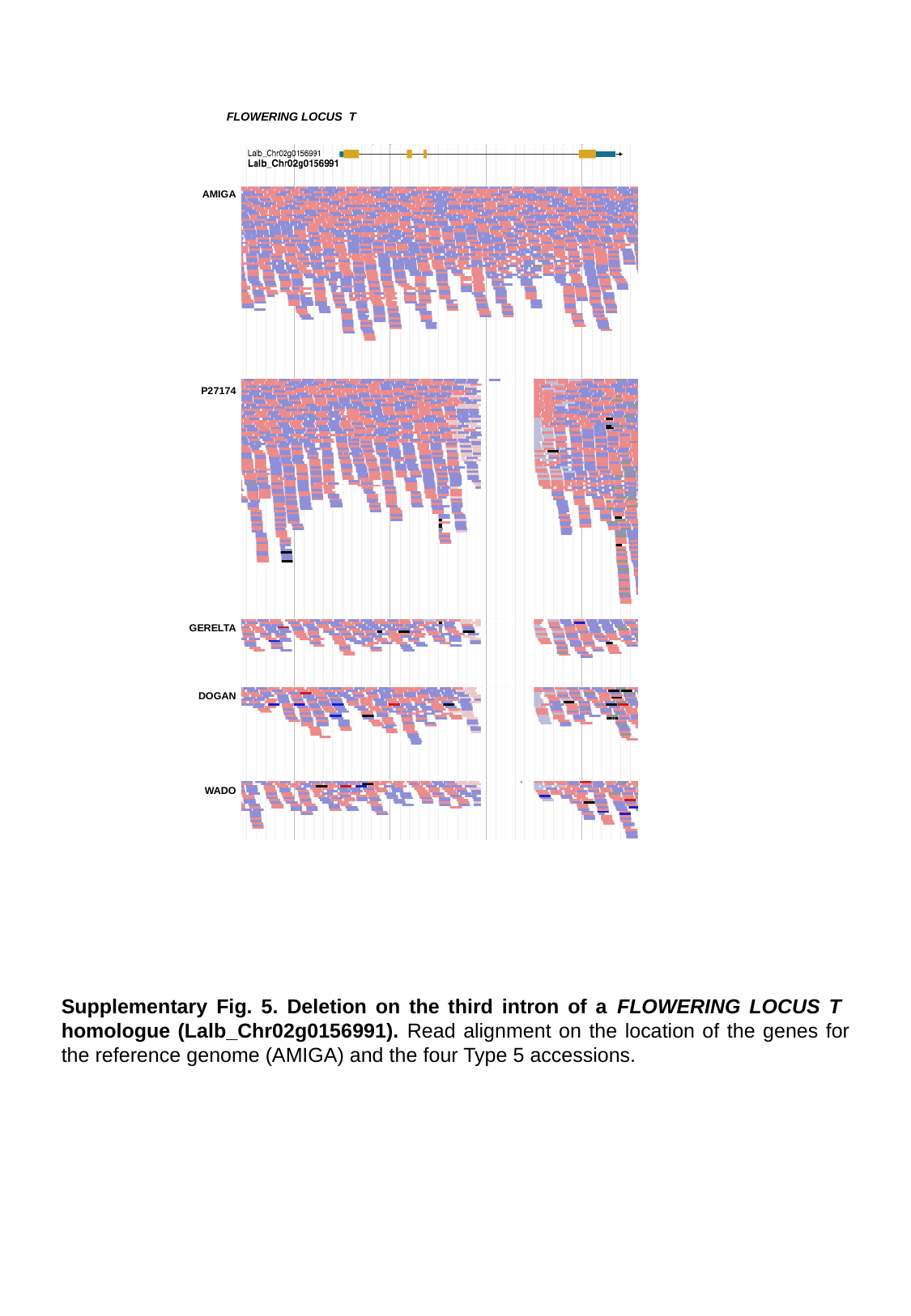

FLOWERING LOCUS T
AMIGA
P27174
GERELTA
DOGAN
WADO
Supplementary Fig. 5. Deletion on the third intron of a FLOWERING LOCUS T homologue (Lalb_Chr02g0156991). Read alignment on the location of the genes for the reference genome (AMIGA) and the four Type 5 accessions.
