## Supplementary Tables for "Pangenome of white lupin provides insights into the diversity of the species"

**Supplementary Table 1. Accessions of white lupin used in the pangenome construction.**

| **Accession** | **Original country** | **Domestication status** |
| --- | --- | --- |
| ALB01 | Algeria | Landrace |
| Amiga | Chile | Cultivar |
| Badajoz | Spain | Landrace |
| Batsi_Wild | Greece | Wild |
| Clovis | France | Cultivar |
| Dieta | U.K. | Cultivar |
| Dogan | Ethiopia | Landrace |
| EGY6484B | Egypt | Landrace |
| Energy | France | Cultivar |
| Feodora | Germany | Cultivar |
| Figaro | France | Cultivar |
| Gerelta | Ethiopia | Landrace |
| GR38-Megalopolis | Greece | Wild |
| Graecus | Hungary | Wild |
| GRC5262B | Greece | Wild |
| Gyulatanya | Hungary | Cultivar |
| Hansa | Germany | Cultivar |
| Kalina | Poland | Cultivar |
| Kiev Mutant | Ukraine | Cultivar |
| Lucky | France | Cultivar |
| Luxe | France | Cultivar |
| Magnus | France | Cultivar |
| Murringo | Australia | Cultivar |
| N3507 | Israel | Landrace |
| Nahrquell | Germany | Cultivar |
| Neuland | Germany | Cultivar |
| Neutra | Germany | Cultivar |
| Orus | France | Cultivar |
| P21525 | Chile | Cultivar |
| P27174 | Ethiopia | Landrace |
| Poutignano | Italy | Landrace |
| Primorsky | Russia | Cultivar |
| Shinfield | U.K. | Cultivar |
| Start | Russia | Cultivar |
| SYR6258B | Syria | Landrace |
| Tombowskij Skorospielyj | Russia | Cultivar |
| Ulysse | France | Cultivar |
| Volodia | Germany | Cultivar |
| Wado | Ethiopia | Landrace |

**Supplementary Table 2. Summary of re-sequencing of white lupin accession using short reads produced in this report.**

| **Acession** | **Number of reads** | **Sequencing depth (x)** |
| --- | --- | --- |
| P21525 | 50 445 994 | 17.5974 |
| ALB01 | 62 031 024 | 21.6387 |
| Badajoz | 52 323 252 | 18.2523 |
| Batsi_Wild | 72 214 902 | 25.1912 |
| Dogan | 62 163 470 | 21.6849 |
| EGY6484 | 49 371 940 | 17.2228 |
| Gerelta-2 | 47 922 500 | 16.7172 |
| GRC5262B | 69 488 018 | 24.24 |
| Gyulatanya | 72 232 012 | 25.1972 |
| Hansa | 49 540 834 | 17.2817 |
| Kalina | 47 682 910 | 16.6336 |
| Murringo | 46 095 750 | 16.0799 |
| N3507 | 59 689 068 | 20.8218 |
| Nahrquell | 50 061 858 | 17.4634 |
| Neuland | 52 169 244 | 18.1986 |
| Neutra | 54 662 092 | 19.0682 |
| Poutignano | 36 943 998 | 12.8874 |
| Primorsky | 60 568 318 | 21.1285 |
| Shinfield | 51 076 124 | 17.8173 |
| Start | 55 437 782 | 19.3388 |
| SYR6258B | 62 770 940 | 21.8968 |
| Tombowskij S. | 40 588 546 | 14.1588 |
| Volodia | 49 050 310 | 17.1106 |
| Wado | 57 808 232 | 20.1657 |
